# Frequent recombination of pneumococcal capsule highlights future risks of emergence of novel serotypes

**DOI:** 10.1101/098335

**Authors:** Rafał J. Mostowy, Nicholas J. Croucher, Nicola De Maio, Claire Chewapreecha, Susannah J. Salter, Paul Turner, David M. Aanensen, Stephen D. Bentley, Xavier Didelot, Christophe Fraser

## Abstract

Capsular diversity of *Streptococcus pneumoniae* constitutes a major obstacle in eliminating the pneumococcal disease. Such diversity is genetically encoded by almost 100 variants of the capsule polysaccharide locus (*cps*). However, the evolutionary dynamics of the capsule – the target of the currently used vaccines – remains not fully understood. Here, using genetic data from 4,469 bacterial isolates, we found *cps* to be an evolutionary hotspot with elevated substitution and recombination rates. These rates were a consequence of altered selection at this locus, supporting the hypothesis that the capsule has an increased potential to generate novel diversity compared to the rest of the genome. Analysis of twelve serogroups revealed their complex evolutionary history, which was principally driven by recombination with other serogroups and other streptococci. We observed significant variation in recombination rates between different serogroups. This variation could only be partially explained by the lineage-specific recombination rate, the remaining factors being likely driven by serogroup-specific ecology and epidemiology. Finally, we discovered two previously unobserved mosaic serotypes in the densely sampled collection from Mae La, Thailand, here termed 10X and 21X. Our results thus emphasise the strong adaptive potential of the bacterium by its ability to generate novel serotypes by recombination.

## Introduction

*Streptococcus pneumoniae* is a human bacterial commensal and pathogen, estimated to be the cause of death in over 500,000 children under 5 years of age each year worldwide (WHO, 2012). The bacterium’s capacity to cause disease is associated with its possession of several virulence factors, of which the most important is the surface polysaccharide capsule (Briles *et al*., 1992; Morona *et al*., 2004; Kadioglu *et al*., 2008; Hyams *et al*., 2010). The polysaccharide diversity constitutes a major challenge for eliminating pneumococcal disease. Being exposed to antibody binding as the outermost layer of the bacterium, the capsule is the target of all licensed pneumococcal vaccines. The most commonly used conjugate vaccines currently target eleven or thirteen of the most common capsular types (serotypes). However, as of today almost 100 distinct serotypes have been described and recognised. Each serotype has a unique, experimentally confirmed serological profile (Henrichsen, 1995), and for many of them the biochemical structure is known (Geno *et al*., 2015).

Systematic genetic sequencing revealed that the diversity of the capsule locus, *cps*, alone forms a repertoire of almost 2,000 unique genes (Bentley *et al*., 2006). These genes are divided based on their functions and form three major groups (Yother, 2011; Geno *et al*., 2015). The first group is located upstream of the locus and consists of modulatory cpsABCD genes (*wzg*, *wzh*, *wzd*, *wze*), which are common to almost all serotypes. The second group are serotype-specific genes (i.e., glycosyltrasferases) with polymer-specific functions, and these define a serotype. Finally, many serotypes carry sugar-synthesis genes needed for capsule production (e.g., rhamnose genes). Comparison of the genetic content of different serotypes demonstrated that capsular-gene acquisition and loss had been the underlying cause of emergence of many serotypes (Aanensen *et al*., 2007; Mavroidi *et al*., 2007). This is not surprising as the pneumococcus is known to undergo frequent recombination (Feil *et al*., 2000; Henriques-Normark *et al*., 2008; Vos and Didelot, 2009), and the capsular locus was shown to be a recombination hotspot (Croucher *et al*., 2011; Chewapreecha *et al*., 2014). Furthermore, we know from previous studies that the extent of within-serotype diversity is under-appreciated, with many hybrid serotypes circulating in the population (Salter *et al*., 2012; van Tonder *et al*., 2016). However, the evolutionary dynamics, and hence the full adaptive potential of pneumococcal capsular polysaccharides, are not well understood.

The aim of this study was to gain a high-resolution view of the evolution of capsular polysaccharides in *S. pneumoniae*. In particular, we wanted to infer the rates of evolution and recombination within the capsular locus, compare these parameters between different serogroups, and compare the relationship between evolution affecting capsular genes and that affecting the remainder of the genome. To this end, we analysed capsular diversity in a collection of 4,469 bacterial isolates form several different studies. Our approach allowed us to observe the evolution of the pneumococcus from the point of view of the capsule itself, subdivided in major serotypes and serogroups, with the tree showing the evolution of the capsular locus and the tips of the tree containing the information about how the capsule changes between different genomic backgrounds. By disentangling horizontal from vertical genetic changes, we gained insight into the timescales of diversification and recombination in capsular genes. This approach brings novel qualitative and quantitative insight into the evolution of serotypes, the principal target of current vaccines.

## Results

### Species-wide serotype diversity

To study the evolution of the capsular locus, we analysed several collections of pneumococcal isolates including two large carriage cohorts from MaeLa, Thailand (Chewapreecha *et al*., 2014) and Massachusetts (Croucher *et al*., 2013); three widespread lineages, PMEN1, PMEN2 and PMEN14 (Croucher *et al*., 2011, 2014b,a); capsular reference collection (Bentley *et al*., 2006); Dutch isolates from invasive disease (Elberse *et al*., 2011); and publically available reference genomes the European Nucleotide Archive (ENA). This gave a total number of 4,469 isolates from 29 countries and 5 continents, as shown in Figure 1A. Illumina-sequenced isolates (96%) were reassembled using a novel pipeline (see Text S1). In total, we obtained 3,813 full capsular sequences, which were serotyped *in silico* (see Methods and Table S1). Figure 1B shows the observed serotype distribution, with 47% of the identified capsular sequences being serotypes targeted by the seven-valent pneumoccal conjugate vaccine (PCV7) and 59% being serotypes targeted by the more recent thirteen-valent vaccine (PCV13). Altogether we identified 96 reference serotypes consisting of 254 glycosyltransferase gene families (henceforth referred to as capsular-specific genes), amongst which we discovered two previously unseen, putative serotypes, here referred to as serotype 10X and serotype 21X, respectively (see Figures S1/S2). Putative serotype 10X was found in five isolates from MaeLa and originally classified as 10B, 10F or 33B. Accordingly, the genetic structure suggests that it is a mosaic of 10C or 10F with another serotype, possibly 35B. Putative serotype 21X was found in three isolates from MaeLa, and its genetic structure suggests that it arose as a recombination between 6C/6D and 39. Indeed, 21X reacted to antiserum which covers serotypes 33 and 39, and the presence of the novel, functional polysaccharide was reconfirmed (see Text S1).

**Figure 1:**
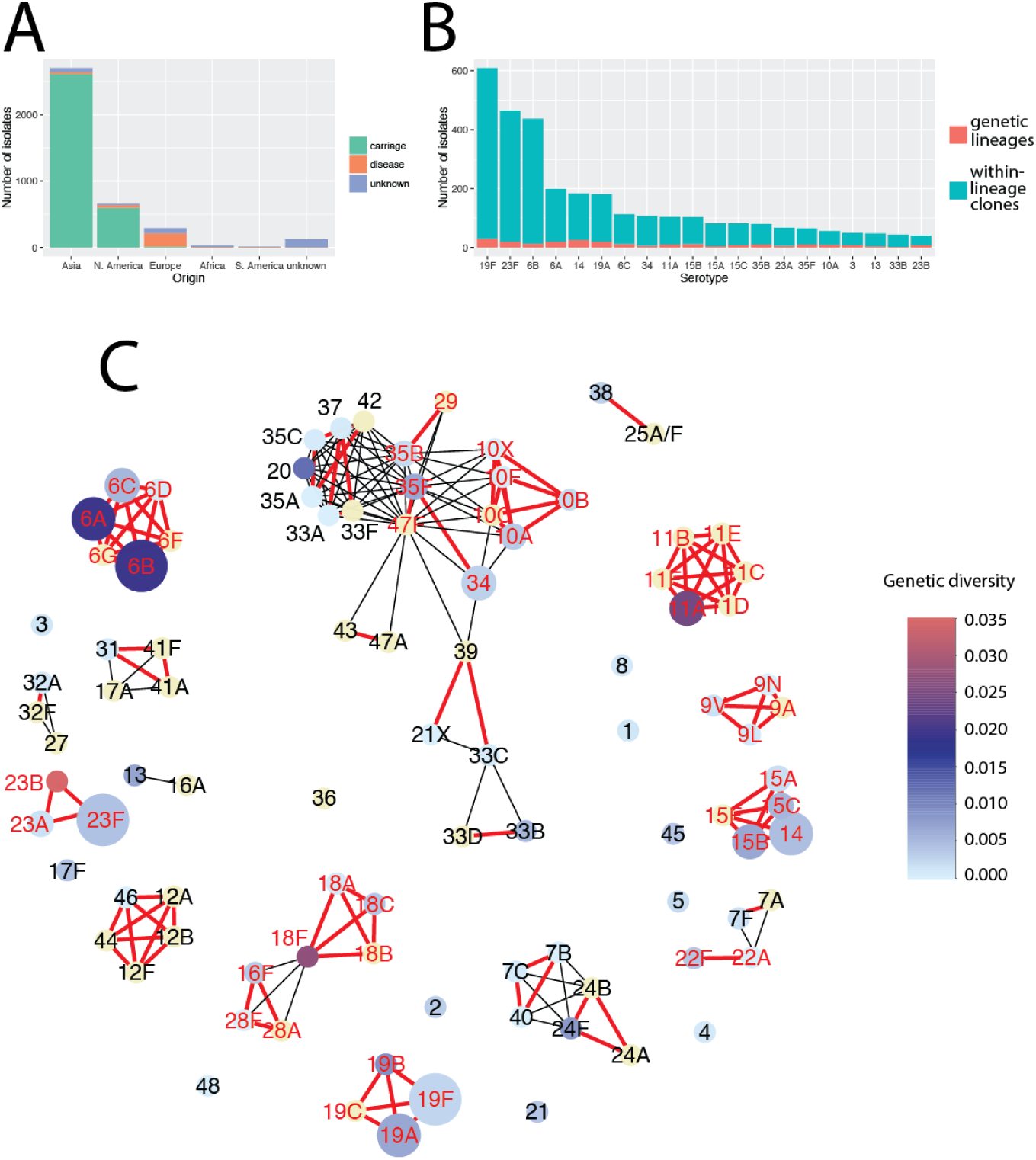
Properties of the dataset. (A) The distribution of the number of isolates stratified by geographical location and the source of isolation (carriage, disease or unknown). (B) The distribution of the number of serotypes. Red bars show the number of genetic serotype lineages, defined by clusters of sequences which can be connected by linking all pairs which differ by two substitutions or less. Blue bars show the remaining count such that the total height equals the full sample size. (C) A diversity network, where each node is represented by a reference sequence and an edge links two nodes if they are similar, i.e., share a minimum proportion χ of the homologies. Red, bold edges show the conservative network where the minimum similarity was defined as sharing χ = 0.58 of homologies. Black edges show the additional connections obtained in a liberal network, where the minimum similarity was defined as sharing χ = 0.36 of the homologies. The size of each node reflects the full sample size, and the colour shows the within-serotype capsular genetic diversity for all non-identical isolates measured using the mean pairwise Kimura K80 distance (full diversity distribution is shown in Figure S4). Red labels of serotype nodes denote genetic serogroups which are analysed in detail below.

To visualise capsular diversity within the pneumococcus, we generated a network with nodes represented by reference sequences and edges linking serotypes with similar capsule-specific gene content. Fig. 1C shows this similarity network for two different thresholds, conservative and liberal, with two corresponding edge types. We see that serotype clusters in the conservative network (henceforth referred to as genetic serogroups) are often congruent with phenotypic serogroups. Nevertheless, there are exceptions: for example serotype 16F clusters with serogroup 28 with conservative threshold, but does not cluster with 16A even with liberal threshold. These observations are consistent with earlier findings and highlight the complexity of the polysaccharide genotype-phenotype map (Aanensen *et al*., 2007; Mavroidi *et al*., 2007). This approach also allows identification of mosaic triplets (nodes connecting other groups), which denote potential introgressive descents (Bapteste *et al*., 2012). In this way one can quickly identify some of the mosaic serotypes, for example 18F (shares *wcxM* gene with 18A/18B/18C and 28A/28F/16F) or 22A (shares *wcwC* gene with 22F and 7A/7F). Furthermore, the largest connected component, which includes serogroups 10, 33, 34, 35 and others, is the most mosaic group of serotypes. Indeed, a closer look suggests that its members consist of the most interconnected serotypes (see Fig. S3). Therefore, we can conclude that many serotypes are highly mosaic in nature and their evolution was likely driven by horizontal transfer of DNA.

### Recombination drives emergence of serotypes

We next investigated the diversification of different serogroups into serotypes. To this end, we focused on the genetic serogroups defined by red edges in Fig. 1C (nodes marked with red labels). As these groups share a large majority of their genetic content, the traditional population genetic methods can be applied to infer the basic evolutionary parameters. We analysed twelve most numerous and diverse serogroups, here referred to as serogroups: 6, 19, 23, 14/15, 18, 10, 11, 9, 34/35, 16/28, 29/35 and 22 (see Fig. S4 for diversity distribution). In brief, sequences from each group were aligned and the population genetic structure was analysed, with recombinations within each sub-population identified using two different methods (Falush *et al*., 2003; Croucher *et al*., 2015a). The recombinant fragments were then removed from the alignment and the resulting clonal alignment was used to construct a tree of the serogroup, with recombinations mapped onto the tree. Figure 2 summarises the inferred model of evolution for the four most common serogroups, 6, 19, 23 and 14/15. The details of the analysis for all serogroups are given in Supplementary Text S1 and Figs S15-S38. Here we briefly summarise the main findings.

**Figure 2:**
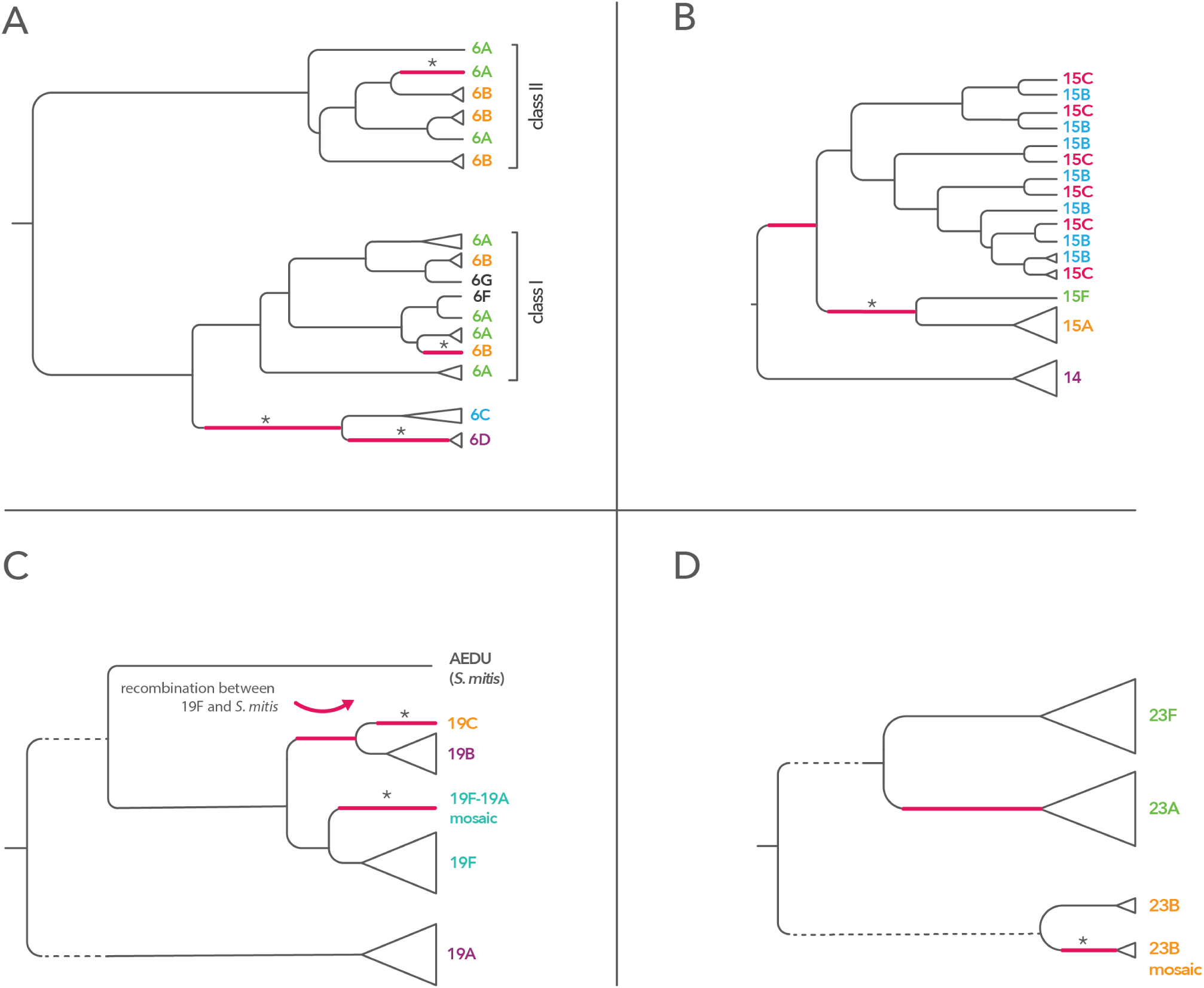
Evolution of the four most common serogroups. Schematic dendogram based on the inferred clonal trees is shown for the four most common serogroups in this study, serogroup 6 (panel A), serogroup 14/15 (panel B), serogroup 19 (panel C), and serogroup 23 (panel D). Branches where recombinations occurred are coloured red, with a star sign marking those branches where there was statistical support for the recombination (using Structure or Gubbins). Clonal uncertainty due to the suggested model is reflected by dashed branches.

First, the critical event in the emergence of at least seven serotypes was a recombination importing extensive genetic diversity, as indicated by the causative change being associated with a cluster of polymorphisms on the ancestral branches of these serotypes. In four of these cases, the source could be identified (see Table 1 and Text S1). Second, in serogroup 19 we hypothesise that the 19B/19C clade arose by arose by recombination of 19F with *S. mitis*, and in serogroup 23 that serotype 23A emerged as a recombination of 23F and a capsule of an unknown source. Third, we detected many recombinations which did not change serotype but sometimes produced mosaic isolates, for example 19A/19F mosaic, 6B-I/6B-II mosaic or 23B-mosaic. Fourth, population genetic structure of the common *wz*-genes shows presence of many older, undetected recombinations (see Fig. S5). Finally, in many cases we observed that a simple model of gene gain and loss cannot explain the observed patterns of diversity. In particular, we found serotypes 6A/6B, 15B/15C and 18B/18C to have emerged on multiple independent occasions, some of which were due to homologous recombination (though in serogroup 15 it could be due to intragenomic recombinations; see Fig. 2 and Text S1). Hence the evolutionary history of these serotypes is a complex story of repeated recombinations of differing phenotypic consequences.

**Table 1:** Direct evidence for emergence of serotypes by recombination. The table summarises the cases in recombinations directly detected in our approach were associated with the emergence of new serotypes. In some cases the direction of recombination could not be established; in this case multiple serotypes were given in the second column. Details of the analysis are given in Text S1.

| serogroup | serotype(s) | gene(s) affected | likely source |
| --- | --- | --- | --- |
| 6 | 6C | *wciN* | unknown (homolog in 21X) |
| 6 | 6D | *wciP*, *wzy*, *wzx* | 6B-II |
| 19 | 19B/19C | *wchU* | *S. mitis* |
| 10 | 10C | *wcrW* | 10A |
| 34/35 | 47F | *whaI* | 47A |
| 16/28 | 28A | *wciU* | unknown |
| 22 | 22A/22F | *wcwC* | unknown |

### Molecular clock of the capsule

We next wanted to learn about the timescales of the evolutionary process within the capsule.Unfortunately, the serogroup alignments did not have enough temporal signal to robustly infer the substitution rate of the capsular locus. Therefore, we used whole-genome collections of three lineages PMEN1, PMEN2 and PMEN14, respectively (Croucher *et al*., 2011, 2014b,a). We inferred alignments with recombinations removed using the software Gubbins (Croucher *et al*., 2015a). For each lineage, we simultaneously fitted two separate clock models, one to the entire alignment with the capsule removed, and the other one to the capsule only, defined by coordinates of *dexB* and *aliA* genes (see Methods), all using BEAST2 (Bouckaert *et al*., 2014). Results are displayed in Figure 3A and show that in all three lineages we observed a roughly 2.5 times higher clock rate at the capsular locus than in the rest of the genome. The distribution of SNPs across capsular genes (see Fig. S6) suggests no bias at transposable elements, which can sometimes produce false-positive substitutions due to their repetitive nature. We can thus conclude that the observed substitution rate is not an artefact of data assembly. To investigate whether the increased substitution rate at the capsule is due to varying rates of selection between different proteins, we estimated the dN/dS ratio using the CODEML approach (Yang, 2007) for the capsule versus all other genes (see Methods). Results are shown in Figure 3B and suggest that the capsule has a higher proportion of non-synonymous substitutions compared to the rest of the genome. It is unclear whether the increased dN/dS is due to positive, diversifying selection or relaxed purifying selection.

**Figure 3:**
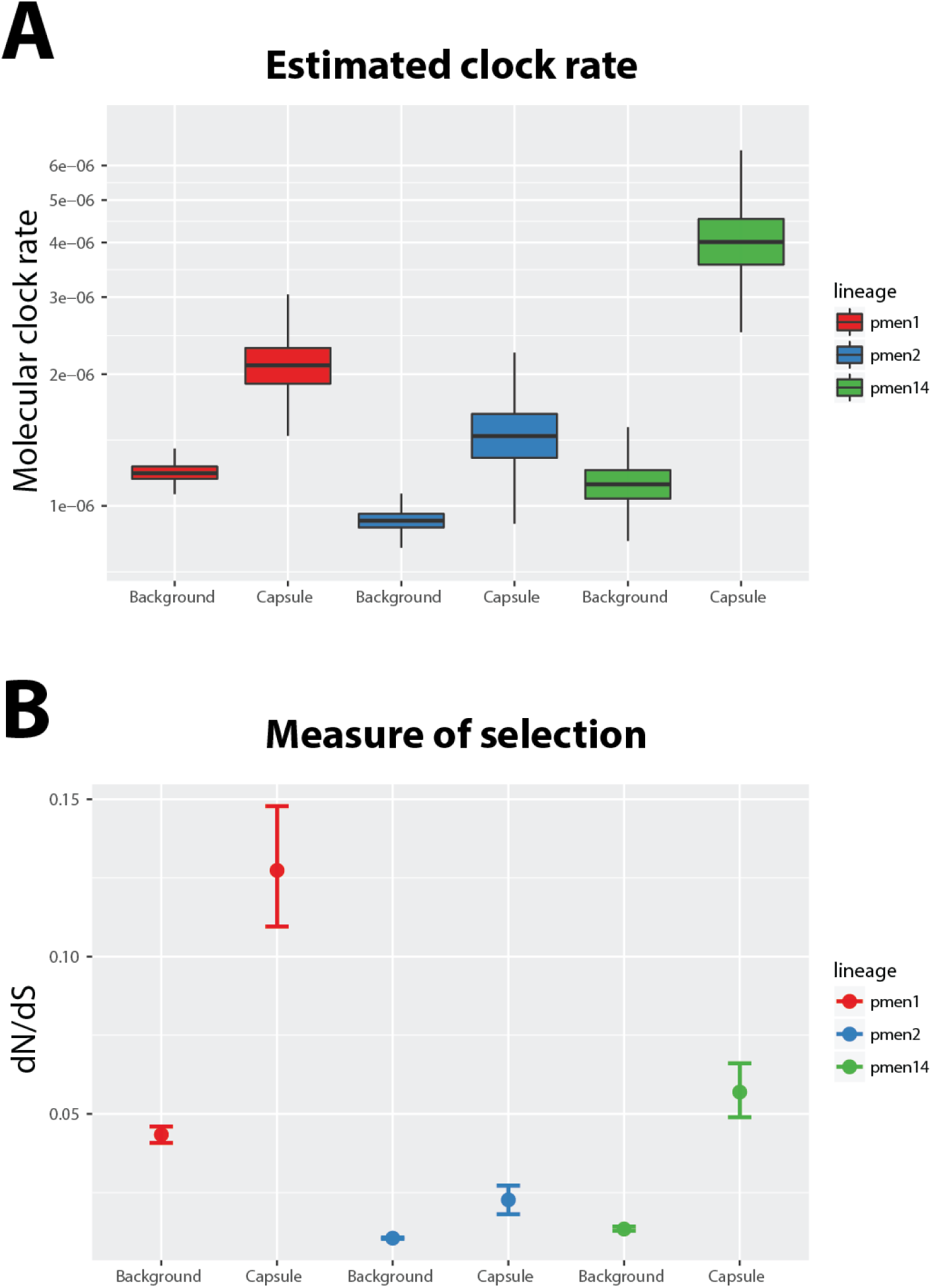
The molecular clock rate of the capsular locus versus background in three different lineages, PMEN1, PMEN2 and PMEN14. (A) The bar plots show the inferred molecular clock rate of the whole-genome alignment with the capsule removed (“Background”), and to the capsule-only alignment (“Capsule”). (B) Comparison of the selective pressures exerted on the background vs. capsule as inferred by dN/dS.

To examine whether there may have been any undetected recombinations in the capsule, which in turn could explain the elevated substitution rate, we tested for the levels of homoplasies using `phitest` (Bruen *et al*., 2006). In all three cases we found non-significant results (p = 0.1, p = 1 and p = 0.1, for PMEN1, PMEN2 and PMEN14, respectively). A closer examination also does not point to a substantial impact of undetected recombinations. In PMEN1 and PMEN2 the proportion of homoplasic substitutions was only 6% and 7%, respectively, and in PMEN2 we would not expect a substantial impact as many representatives of this clone had their competence turned off by a phage insertion. In PMEN14 the proportion of homoplasic substitutions was 21%, but in this case the higher number is likely due to selection: of 48 independently occurring SNPs across all capsular coding regions, 38 were located within two genes, *wzd* and *wze* (see Fig. S5). The estimated dN/dS in those two genes was many times higher than across all capsule: 1.25 (95% CIs: 0.70-2.65) compared to 0.057 (0.049-0.066). Thus, even though the impact of recombination on the elevated substitution rate cannot be excluded, the higher molecular clock observed at the capsule is most likely explained by an impact of selection.

### Variability in recombination rates between serogroups

Using the capsular clock rate estimated for PMEN collections and the sampling dates obtained for the isolates used in this study, we next estimated the branch lengths of the ML trees for each of the serogroups. This also allowed us to infer the divergence times of the capsular clonal tree (shown in Figs. S7-S10) with the times of occurrence of corresponding recombination events, as well as to estimate the recombination rate for each serogroup in more intuitive units. Figure 4A shows the comparison of the obtained recombination rate estimates for the 12 examined serogroups. The results demonstrate a significant heterogeneity in recombination rates across all serogroups (oneway anova, p < 10^−16^), though pairwise differences were often statistically non-significant. To test whether sampling can explain the observed variance in recombination rates, we examined associations between the rates and five measures of sampling: number of isolates, number of nonidentical isolates, clonal diversity, number of countries in which each serogroup was sampled and number of clonal complexes (see Table S1 in Text S1). We did not find a significant association between the recombination rate and any of the quantities (Spearman rank test, p > 0.05 in all cases). As corresponds to theoretical expectations (see Fig. S11), we conclude that sampling cannot explain the observed variation in recombination rates.

**Figure 4:**
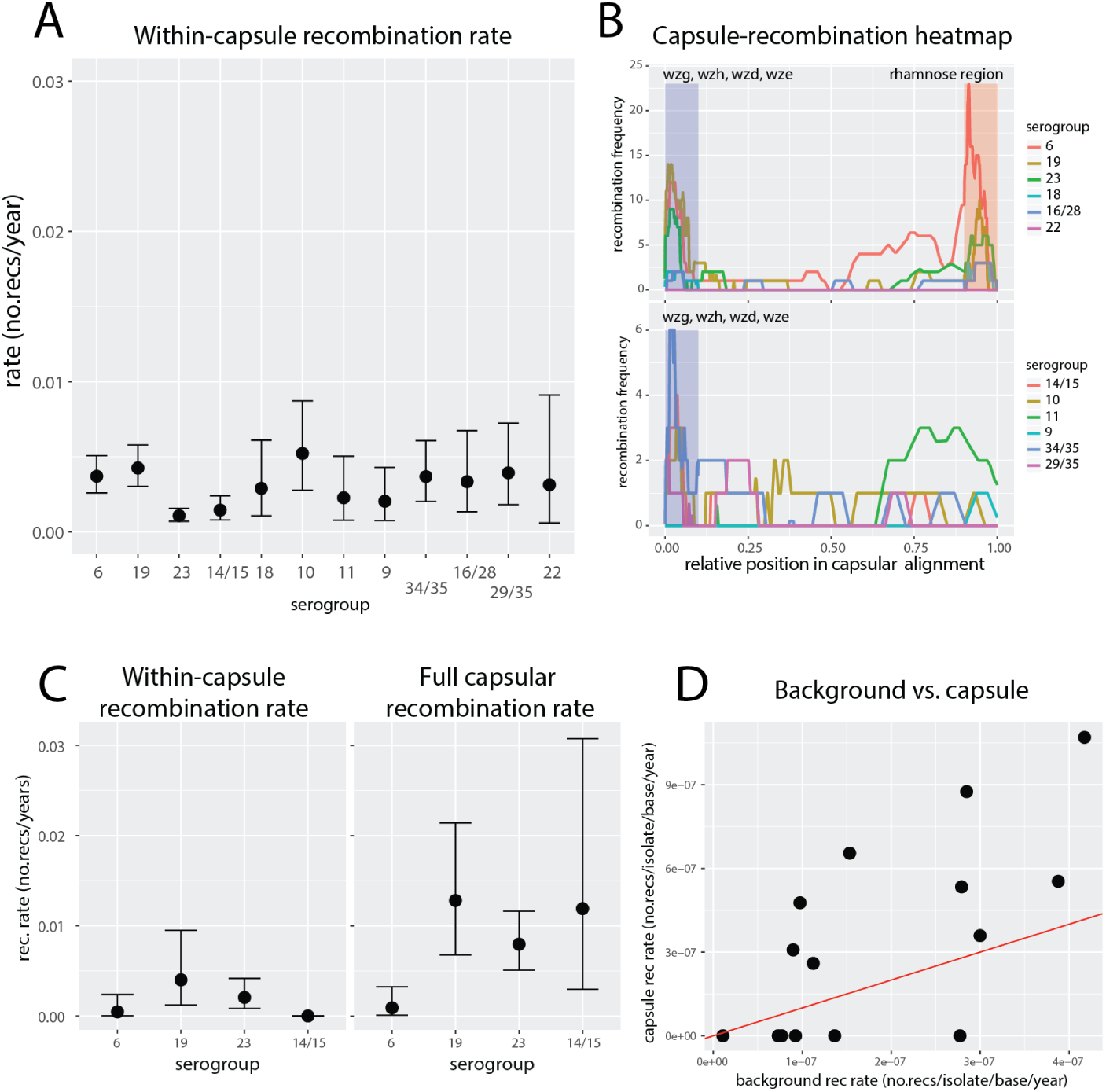
Recombination rates within the capsular locus. (A) Recombination rates estimated for the 12 serogroups used in this study, with the relevant 95% confidence intervals. By definition these rates do not include long, capsule-spanning recombinations which are invisible from the point of view of capsular alignment. (B) Recombination heatmap at the capsular locus, with different colours showing number of recombinations for different serogroups, with rhamnose genes (top) and without rhamnose genes (bottom). Recombination positions were normalised such that total alignment length was 1 in all serogroups. Additionally, for the sake of comparison, the upstream *wg*-region (blue) and the rhamnose region (red) were normalised to 10% of the length each. (C) Recombination frequency measured using whole-genome approach, with capsule-contained events (left) vs. all capsule events (right). (D) Recombination rate at the genomic background (excluding events at the capsule) versus recombination rate of events affecting the capsule, estimated from whole-genome alignments. The rates were normalised per base using mean alignment lengths of whole-genome and capsule, respectively. The y = χ line is shown in red.

The second hypothesis which could explain the variation in recombination rates is the selective pressure to preserve the polysaccharide structure by maintaining capsular genetics. That such selection might act is already suggested by the recombination heatmap across the capsular locus (Fig. 4B): we found the recombination frequency higher at the flanks of the capsule, namely within those genes which are common to many serotypes. One potential explanation for this is that those regions are less likely to disrupt the capsule-specific polysaccharide structure and are thus observed more often due to negative purifying selection. However, more frequently observed recombinations are also expected to occur at a higher rate at relatively conserved sites due to a higher number of possible donor-recipient pairs in the homologous recombination process, which requires sequence-similarity at flanking regions. To circumvent this problem, we used a whole-genome lineage-by-lineage approach in all used data collections to detect recombinations with breakpoints outside the analysed capsular region and thus invisible in our alignments (see Methods). By analysing only lineages which were at least 95% of a single serogroup, we could directly compare the frequency of (i) non-capsular homologous recombinations, (ii) all recombinations affecting the capsule and (iii) recombinations contained within the capsule. Fig. 4C shows the comparison of the mean within-capsule recombination rate (left) and the mean total capsule recombination rate (right). Under a neutral model we would expect 49.6% of all capsular recombinations to be contained within the capsular locus (see Text S1 for full calculation). Comparison of the recombination rate ratios revealed that only serogroup 6 was found to lie within that range (0.500, CI 0.136 – 0.733), while for the other three serogroups we found less within-capsule recombinations than expected (serogroup 19: 0.313, CI 0.177 – 0.443; serogroup 23: 0.258, CI 0.161 – 0.357; serogroup 14/15: 0). These results suggest that within-capsular recombinations may be under different selective pressures in different serogroups (see Discussion).

Another hypothesis that could explain the variation in recombination rates is the background recombination rate, which is known to vary between lineages (Croucher *et al*., 2013). To investigate this we asked whether the genomic recombination rate is a good predictor of the capsule recombination rate (Fig. 4D). We found that the former explains roughly half of the variance in recombination rates between serotypes (linear model, R^2^ = 0.47, p < 0.01). Finally, we found no significant relation between the mean serogroup capsule size, as measured experimentally (Weinberger *et al*., 2009), and the estimated recombination rate (see Fig. S12). As capsules are known to constitute a physical barrier to incoming transformation events (Schaffner *et al*., 2014), this result suggests that the variation in observed recombination rates cannot be explained by raw transformation rates, but rather reflects the differential effect of selection on the capsular locus.

### Origin of capsular recombinations

We next investigated the origin of the identified recombination events. To this end, we used BLAST to identify close hits (defined by min. 90% identity; see Methods) with multiple hits assigned a proportionally lower weight; otherwise the origin was considered unknown. We also included a set of 50 *S. mitis* sequences mentioned in Table S1. Fig. 5A shows a recombination flow diagram, namely a directed network with arrows indicating the direction of recombination between different serogroups. We identified potential source for 91% of recombinations. It is unclear whether the remaining recombinations descended from the same or other bacterial species. However, we would not expect to find many cases of inter-species recombinations for several different reasons, including biological ones (stronger purifying selection of more diverse imports) and methodological ones (under-sequenced diversity of non-pneumococcal streptococci). Nevertheless, in the case of serogroup 19 we found four recombinations with close homology to *S. mitis*, and in serotype 14 one recombination with close homology to *S. oralis*. We also found that more recombinations originated in other serogroups compared to same serogroup (156 vs. 105), consistent with the observation in that most recombinations are found in genes common in other serotypes (cf. Fig. 4C). To explore this dependence for each serogroup, we quantified the number of recombinations originating in the same (“self”) versus in different (“non-self”) serogroup (Figure 5B). We found that most serogroups have more non-self recombinations, but in three serogroups (6, 10 and 11) the majority of recombinations originated in the same serogroup. (The results were qualitatively similar when considering only capsule-specific recombinations.) These results could explain a lower within-capsule recombination rate in serogroups 19, 23 and 14/15 than expected: recombinations originating in other serogroups are more likely to disrupt the capsule, and hence be selected against.

**Figure 5:**
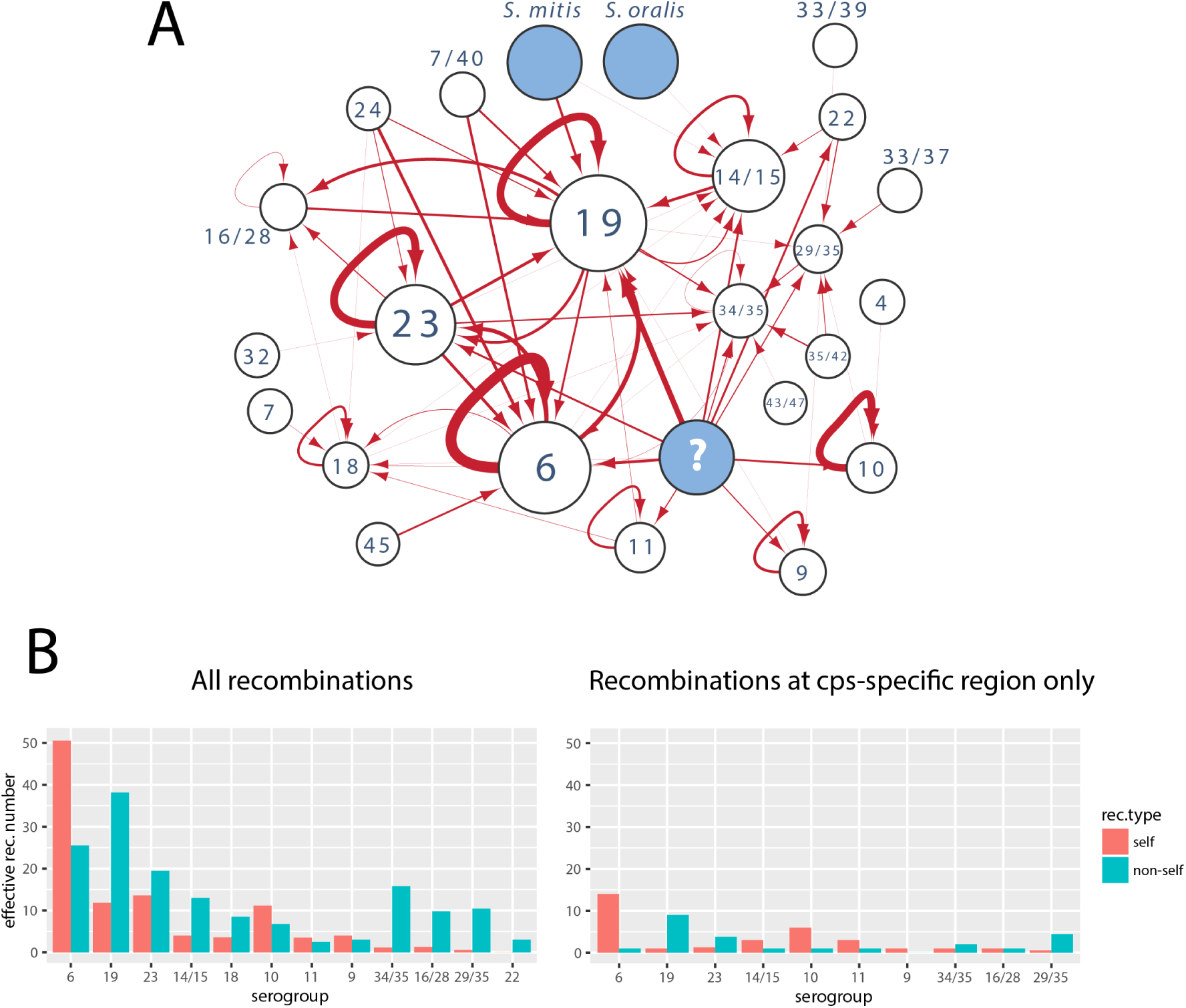
Origin of identified recombinations. (A) The network shows the recombination flow among the serogroups defined in Figure 1A: nodes correspond to serogroups and arrows correspond to the direction of recombination flow based on the most likely origin of the putative recombination events. The width of arrows reflects the number of recombination events (between 0 and 51) and the size of the nodes reflects the number of isolates within the serogroup (except for ‘unknown’ and other streptococci). (B) Proportion of recombination events originating in the same serogroup (self) versus another serogroup (non-self) for each serogroup. The plot on the left shows all recombinations, while the plot on the right shows recombinations which overlap with the capsule-specific region (minimum 100bp overlap).

### Capsular lineage jumping

Analysis of capsular sequences allows the evolution of the species to be observed from the point of view of the antigen which occasionally alters its genetic background via ‘lineage jumping’. Such jumps should be observed as alterations of clonal complexes (lineages) within individual clades of serotype trees. The example of serogroup 6 (Fig. 6A) suggests a large within-clade variation of clonal complexes. To estimate the rates of lineage-jumping for four major serogroups (6, 19, 23 and 14/15), we defined a clonal complex (CC) using eBurst (Feil *et al*., 2004) by linking isolates with 6/7 MLST-locus identity, and a clonal complex group (CCG) by linking isolates with 5/7 MLST-locus identity. We next used BEAST2 to predict the rate with which an isolate in each serogroup is expected to jump lineage (see Methods). The results are shown in Fig. 6B. We found the mean jumping-rate between CCs to be 4.7 × 10^−3^ jumps per isolate per year, and between CCGs to be 5.0 × 10^−4^ jumps per isolate per year. If changes between all pairs of CCs were equally likely, we would expect that 57% of them would alter the CCG. Thus, under a random CC-jump model we would expect the CCG jumping rate to be roughly 0.57 times the CC jumping rate. Instead, we found the CCG jumping rate to be lower than expected. These results suggest that pneumococcal serotypes are less likely to jump lineages if those lineages are very distant. This is consistent with the observation that most pneumococcal serotype switches were previously found to occur within a serogroup (Croucher *et al*., 2015b).

**Figure 6:**
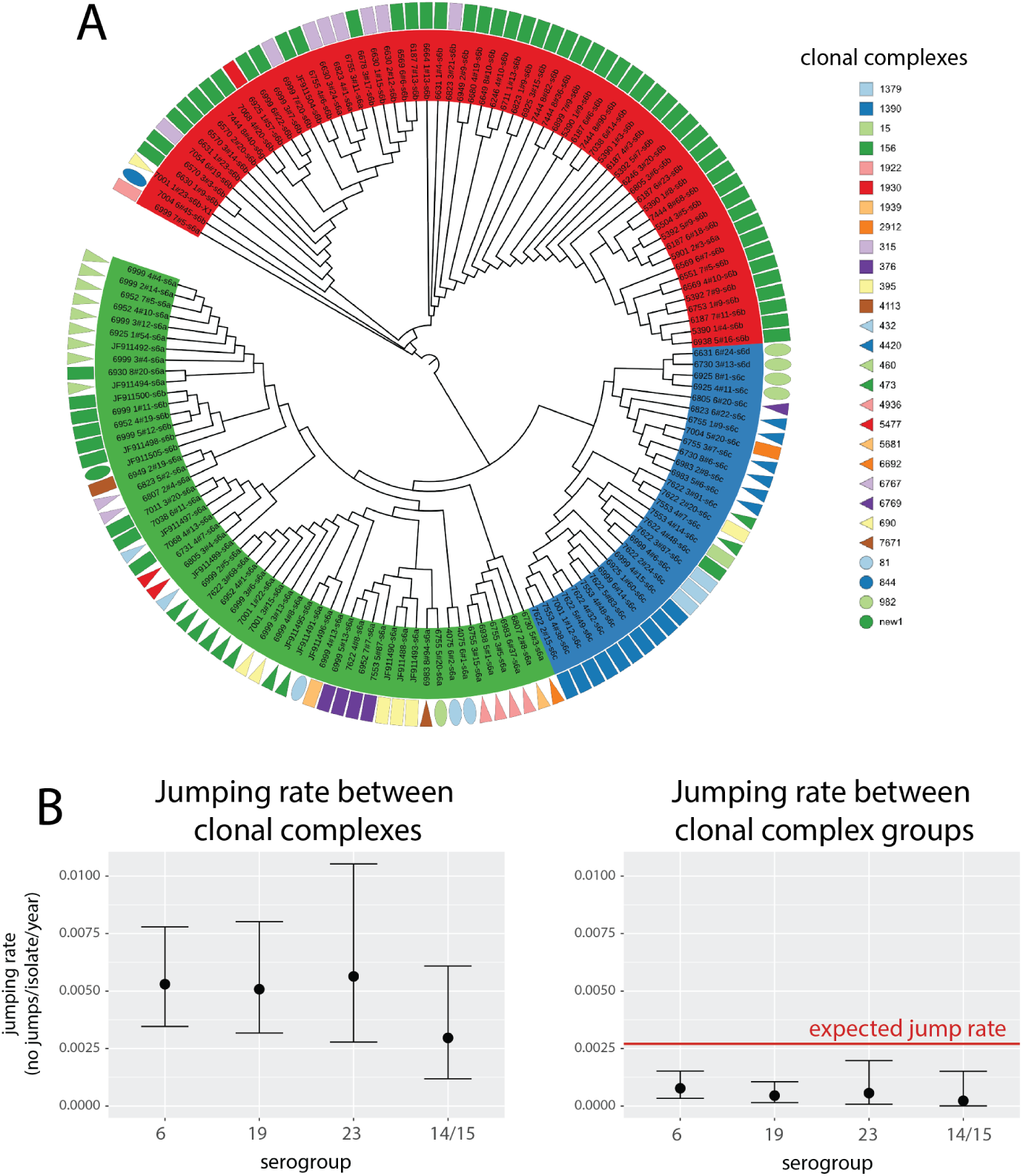
Lineage-jumping dynamics. (A) Dendogram based on the phylogeny of serogroup 6. Tips are grouped according to the three major populations identified: class-I 6A clade (green), 6C/6D clade (blue) and class-II 6B clade (red). Geometric shapes aligned with the tips denote the corresponding clonal complexes of the strains in which the serotype sequence was found. The clades of the tree with branch lengths shorter than 9 × 10^−5^ were collapsed and the most frequent lineage within the clade was plotted. (B) Lineage-jumping rates inferred for the four most frequent serogroups (6, 19, 23 and 14/15) with lineages defined as clonal complexes (CC; left) and as clonal complex groups (CCG; right). The red line shows the CCG jumping rate expected based on the observed CC jumping rate and the assumption that changes between all pairs of CC are equally likely.

## Discussion

The pneumococcal capsule is an evolutionary hotspot, presumably underlying immune selective pressures acting on this major antigen. In the capsule, we found both elevated molecular clock rate and recombination rate compared to the rest of the genome. The first is consistent with previous observations that non-essential bacterial proteins are expected to evolve faster than essential proteins (Jordan *et al*., 2002), and non-encapsulated pneumococci have been known to both persist epidemiologically (Chewapreecha *et al*., 2014) and cause infections (Dixit *et al*., 2016). We also found the recombination rate at the capsular locus to be around 2.5 times higher than in the rest of the genome (see Fig. S13). The simplest explanation is that this is due to increased recombination-detection power due to increased diversity, however recombination also preferentially occurs between closely related isolates (Majewski *et al*., 2000; Majewski, 2001; Ansari and Didelot, 2014). Thus, the recombination rate is more likely elevated for the same reasons as the molecular clock, namely that most of observed recombinations are either those which selection has not yet had time to eliminate, or those promoted by diversifying selection. This hypothesis was in line with two observations. First, the average recombination rate was found to be higher in densely sampled Thai collection than in the US collection (see Fig. S12). We think this is because in densely sampled areas there is a higher chance of finding a rare recombination. Second, recent branches of capsular trees had more recombinations than expected from a neutral model (see Fig. S14). Thus, the resulting relaxed purifying selection produces observed recombination hotspots, which translate into an increased diversity over time on which selection can act.

Interestingly, selective pressure can vary between serogroups. The observation that most recombinations in serogroups 19, 23 14/5 – but not serogroup 6 – originate in other serogroups is in line with the finding that in former three we found less within-capsule recombinations than expected from a neutral model. One potential explanation is that is that the shorter, within-capsule recombinations are under negative selective pressure to preserve the integrity of the capsular locus. The varying selection pressure across serogroups could be due to different epistatic interactions of capsule-specific genes (glycosyltransferases) as well as also because some serotypes are more likely to recombine with serotypes from the same serogroup (Croucher *et al*., 2015b). Overall, we can conclude that the observed variation in recombination rates of different serogroups is a combination of microbiological, ecological, epidemiological and evolutionary processes acting on the pneumococcal capsule.

Our results suggest that inter-species recombination plays an important role in the evolution of the cps locus. One prominent example is the emergence of the 19B/19C clade by recombination of 19F with *Streptococcus mitis*. Furthermore, five detected recombinations bore close resemblance to *S. mitis* and *S. oralis* isolates. In addition, many glycosyltransferase genes are shared between different species in the mitis group (Kilian *et al*., 2014). Given the scale of genetic diversity across the mitis group (Kilian *et al*., 2008), this suggests that many older, hypothesized recombinations within the capsule (for example the emergence of 23A by acquiring *wzy* gene) were probably acquired from other closely related bacteria. However, the timescales of such process remain unclear and require a better characterisation of the polysaccharide diversity in all nasopharyngeal bacteria.

All evidence thus points at *cps* being a genetically plastic and dynamic locus, with recombination being its main evolutionary driver. While most recombinations are expected to either be under weak negative selection or produce non-viable capsules (Park *et al*., 2014), occasionally mosaic, previously unseen capsules can emerge. Indeed, in this study we identified two such mosaic alleles, termed 10X and 21X, while a 33B/33C hybrid was previously identified in an isolate from Denmark (Salter *et al*., 2012). Given (i) that dense sampling leads to the discovery of more recombinations, (ii) that identifying a novel, mosaic capsule requires a detailed, comparative approach, and (iii) the enormous diversity of glycosyltransferases in the microbial world, we can expect that many more such hybrids are circulating around the world. Why are these hybrids not spreading in the population? This could be due to several different factors, including cross-immunity (Lipsitch, 1997), competitive exclusion (Trzcinski *et al*., 2015) or fitness differences (Cobey and Lipsitch, 2012). However, introduction of broader, conjugate vaccines in the future may empty ecological niches occupied by the common serotypes, and provide a selective advantage for some of the rare, mosaic serotypes, which could increase in frequency over time. Therefore, a systematic characterisation of capsular diversity across different nasopharyngeal species is important for a better characterisation of the true pneumococcal adaptive potential

## Materials & Methods

### Isolates

In this study we combined several, previously published genetic and genomic data collections of *Streptococcus pneumoniae*. These include 3,085 isolates from a continuous mother-infant carriage study in the Mae La refugee camp, Thailand (Turner *et al*., 2012; Chewapreecha *et al*., 2014), 616 isolates from children carriage in Massachusetts (Croucher *et al*., 2013), and 605 isolates from the Pneumococcal Molecular Epidemiology Network of three different lineages: PMEN1 (Croucher *et al*., 2011), PMEN2 (Croucher *et al*., 2014b), and PMEN14 (Croucher *et al*., 2014a). We also included 45 sequences of serogroup 6 and 19 isolates from invasive disease from The Netherlands (Elberse *et al*., 2011), 92 reference sequences (Bentley *et al*., 2006; Park *et al*., 2007; Bratcher *et al*., 2011; Oliver *et al*., 2013), and a set of 25 reference genomes of *S. pneumoniae* as found in the European Nucleotide Archive http://www.ebi.ac.uk/genomes/bacteria.html.

### Assembling capsular sequences

All Illumina-sequenced isolates were reassembled using `velvet` (Zerbino and Birney, 2008) with varied k-mer length (between 50% and 90% of the short-read) and the expected coverage (between 5 and 140). The aim was to find an assembly which spanned as much of the entire capsular locus as possible (defined by aligning the assembled and reference sequences using BLASTN with e-value < 10^−50^) in as few contigs as possible; if multiple such assemblies were produced, the one with the least number of N’s and the highest n_50_ value was chosen. Due to the repetitive nature of transposable elements, sequences were assembled without the flanking *dexB*/*aliA* genes and transposable elements. The resulting set of contigs was then analysed for potential misasseblies using `reapr` (Hunt *et al*., 2013) and had gaps filled using `GapFiller` (Nadalin *et al*., 2012). Finally, isolates with the full capsular locus were scaffolded against the corresponding reference sequence using ABACASS (Assefa *et al*., 2009) and GapFiller. All poor quality assemblies were removed from the analysis, leaving 3651 capsular allele sequences as well as 162 previously PCR-sequenced isolates. The list of all isolates used in the study is given in Table S1.

### Capsular diversity analysis

To compare the genetic similarity of different reference serotypes, we first collected a list of all proteins located within the capsular locus of all 96 references used as described previously (Bentley *et al*., 2006; Mavroidi *et al*., 2007). Of those, we only focused on glycosyltransferases thereby excluding overrepresented *wzg*, *wzh*, *wzd*, *wze*, *dexB*, *aliA*, *aliB*, sugar synthesis genes and transposable elements, giving altogether 742 proteins. All protein sequences were then classified into homology groups with a similar approach to (Mavroidi *et al*., 2007). Specifically, all-versus-all `blastp` was run with e-value threshold of 10^−50^ with hits with less than 60% query coverage ignored. The resulting undirected network was analysed with MCL (Enright *et al*., 2002) with inflation value of 2, and the resulting 254 homology groups were identified. A sequence similarity network was constructed using 96 reference serotypes with nodes representing reference isolates and edges representing similar sequences. Similarity was defined as the proportion of shared homologies between any pair; if the score was asymmetric the higher similarity index was taken. A network was built for a given similarity threshold meaning that all pairs above a chosen similarity threshold were connected with an edge. A conservative similarity index of 0.58 produced 40 clusters which were used as a basis for defining a genetic similarity group. Using such clustering, we identified 12 groups which had at least 40 isolates and 500 single nucleotide polymorphisms in the alignment (as given in main text). With the exception of serogroup 19, these genetic similarity groups were identical when defined on the basis significance of shared similarity with threshold of e = 0.01 based on the approach used by Lima-Mendez *et al*. (2008).

### Obtaining clonal trees of serogroups

Genetic serogroups were initially aligned using `progressiveMauve` (Darling *et al*., 2010). The homologous blocks, identified by the command `blastn-task megablast`, were aligned using `mafft` with `-ginsi` option (Katoh *et al*., 2002), while the non-homologous blocks were concatenated to avoid force-alignment. All blocks were then concatenated to produce the full alignment. To infer the population genetic structure of serogroups, the Structure software with linkage model was used (Falush *et al*., 2003). The runs were based on at least 600,000 iterations plus 200,000 burn-in and with multiple chains to insure that the model has converged. The number of populations K was found as the smallest value of K which explained the observed population structure, for which independent runs gave the same output, and which was supported by the value obtained by BAPS (Corander and Marttinen, 2006). In all examined cases, the identified populations corresponded well to major serotypes or serotype groups (see Text S1 for the results of the population structure analyses). Between-population recombinations were defined by Structure as those with the minimum posterior probability for originating in a different population of 0.75 and reaching at least 0.95 at one site, and removed. The initial phylogeny was generated using PhyML (Guindon *et al*., 2010). Next, each population was analysed by Gubbins (Croucher *et al*., 2015a) with the initial phylogeny used as a starting tree. Recombinations identified by Gubbins or Structure were removed. The resulting clonal alignments were then analysed again by Structure to identify potential hierarchical population structure and within-population recombinations which Gubbins could not detect. In the final alignment, all regions identified by Structure or Gubbins were removed from the alignment to generate the final phylogeny. The pattern of recombinations on this tree was predicted by both Gubbins (running a single iteration conditional on the final phylogeny with two window sizes: 1kb and 10kb) and Structure (using the ace() function in ape package in R to predict the most likely ancestral pattern (Paradis *et al*., 2004)). The two types of recombinations were then merged into the final list of recombinations with overlapping blocks merged, however because Gubbins has a more elaborate algorithm of predicting ancestral recombinations based on ancestral SNP reconstruction, Structure-recombinations at internal nodes which did not overlap with Gubbins-recombinations were ignored. All recombinations ancestral to each of the K populations found were ignored due to low detection power of events on long tree branches.

### Estimating timescales using BEAST

To estimate the molecular clock of the capsular locus, three collections of pneumococcal whole-genomes were used, PMEN1, PMEN2 and PMEN14 (Croucher *et al*., 2011, 2014b,a). Each collection was assembled, aligned and recombinations were removed using Gubbins (Croucher *et al*., 2015a), with details of the processing analysis given in the original publications. The alignment was then divided into the capsule-only alignment (defined by the starting position of the *dexB* and the ending position of the *aliA* gene) and the clonal alignment with the capsule removed. The two alignments were then analysed using BEAST2 (Bouckaert *et al*., 2014) using a single analysis with parameters shared between alignments. In particular, we assumed the same substitution model (GTR with four gamma categories), the same tree prior (Coalescent Bayesian Skyline), and tree but different parameters of the clock model (Relaxed Clock Log Normal). We ensured that all parameters were estimated with effective sample size (ESS) above 200. The results of the ucldMean parameter for the capsule were then pooled together to which a log normal distribution was fit, yielding best fit parameter of μ = –12.98. Next, the clonal alignments for each serogroup (i.e., with recombinations removed) were analysed using BEAST2, and the same set of models was fit as in the case of whole genomes with three exceptions. First, we assumed a coalescent constant population tree prior. Second, an informative prior for the molecular (strict) clock was used with the value of μ estimated using whole-genomes. Third, the phylogeny was fixed as the final clonal phylogeny obtained in the previous section, thereby estimating divergence times and the dates for the recombination events assigned to each node of the tree. A similar approach was considered to estimate the lineage-jumping rate. Lineages were identified as clonal complexes (CCs) and clonal complex groups (CCGs; see main text). These lineages were next treated as discrete traits to perform a discrete trait phylogenetic analysis (Lemey *et al*., 2009) by modelling switches of the genomic lineage background as substitutions in the discrete trait. Sampling of isolates was blind to the genomic background, so in this context no sampling bias for the discrete trait analysis was expected (De Maio *et al*., 2015). Each sub-population was considered as a separate phylogeny with the shared genetic clock and the jump-rate shared between phylogenies within the same serogroup. Due to simplicity a homogeneous jumping-rate was assumed.

### Estimation of recombination rates

In order to estimate the recombination rate for each serogroup, we fitted a basic model describing the distribution of recombination events on a tree using a Poisson process; see also Mostowy *et al*. (2014). The number of recombinations at each branch of the tree was modelled as a Poisson-distributed random variable m_i_ with mean λL_i_ where λ is the inferred recombination rate and L_i_ is the branch length in years. The estimated recombination rate λ was the value which maximised the likelihood 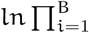 Pois(m_i_; λL_i_), where B is the number of tree branches. A single recombination rate was estimated for each serogroup. To avoid estimate bias we excluded all long branches of the tree, namely branches leading to the most recent common ancestors for each subpopulation (of the K populations estimated by Structure) together with all their ancestor branches.

To estimate recombination rate derived from whole genome data, we used clonal trees obtained as described in the original publications for each of the whole-genome collection used in this work. Briefly, for each bacterial lineage, a clonal tree was estimated with recombinations aligned against nodes of the tree. Genes *dexB* and *aliA* were used to identify the capsule coordinates and distinguish background-from capsule-recombinations. Lineages of predominantly the same serogroup (minimum 95%) were used collectively to estimate the mean recombination rate by combining the information about the number of recombination events on each branch and the corresponding branch length in units of years, and fitting the Poisson model described above. The lineages used to estimate the recombination rates the four analysed serogroups were as follows (please refer to original publications for details). Serogroup 6: Massachusetts (MA) lineages 10, 13 and 14 and PMEN2 (altogether 247 isolates); serogroup 19: MA lineage 15, MaeLa 1 and PMEN14 (altogether 564 isolates); serogroup 23: MA lineage 9 and PMEN1 (altogether 297 isolates); serogroup 14/15: MA lineage 3 and MaeLa lineage 7 (altogether 126 isolates; remaining MaeLa lineages were ignored due to the absence of the full resolution capsular locus in the original alignment).

## Acknowledgements

This work was supported by the EU Marie Skłodowska-Curie Intra-European fellowship (project no. 329515, R-EVOLUTION PNEUMO to R.J.M.), Junior Research Fellowship from Imperial College London (to R.J.M.), Sir Henry Wellcome Fellowship (107376/Z/15/Z to C.C.), NIH MIDAS program (grant U01GM110721 to C.F.) and The Wellcome Trust (grant no. 09805). The authors thank Marc Lipsitch and Bill Hanage for interesting comments and discussions, James McInerney for help with network analyses, Remco Bouckaert and Denise Kühnert for help with BEAST analyses, Aleksandra Królik for help with artwork and Statens Serum Institut for the serotyping work on three 21X isolates.

